# *baal-nf* identifies motif-disrupting variants that decrease transcription factor binding affinity

**DOI:** 10.1101/2025.01.17.633399

**Authors:** Breeshey Roskams-Hieter, Øyvind Almelid, Chris P Ponting

## Abstract

Human traits vary in part due to genetically-determined change of transcription factor (TF) binding affinity within gene regulatory regions. However, few trait-causal variants or mechanisms are known. Here we propose 1,960 variants as strong candidates for causally altering human traits. They were discovered by *baal-nf* which uses ChIP-sequencing data to identify allelic imbalance at heterozygous sites (‘allele-specific binding sites’; ASBs) within affinity-concordant positions within TF- and/or co-factor binding motifs. These variants are evolutionarily conserved, and enriched for trait associations and gene expression QTLs. *baal-nf* and these high-quality ASBs now allow trait variation due to altered TF binding to be investigated.

## Background

Revealing the molecular mechanisms by which DNA variants alter function is critical for understanding how genotypes contribute to physiological traits^1^. DNA variants associated with complex traits and disease mostly lie within non-coding regions, suggesting that trait variation is often caused by genetically altered transcription factor (TF) binding affinity^2,3,4,5,6,7^. Attributing trait variation to a particular TF and its altered site-specific binding affinity, however, remains an unsolved problem in population genetics and functional genomics. Its solution will require accurate and comprehensive catalogues of both TF binding motifs and affinity-altering DNA variants mapped to these motifs^8^.

DNA variants that significantly alter TF binding have been inferred from chromatin immunoprecipitation with sequencing (ChIP-Seq) studies. Rather than comparing across population samples, these experiments use the known genotypes of single cell lines to minimize false variant calls^9,10,11^. TF binding that favours one allele over the other (i.e., allele-specific TF binding sites; ASBs) is inferred at the cell line’s known heterozygous sites when there is a statistically significant imbalance of ChIP-Seq reads mapped to one of the two alleles. When inferring ASBs, however, care is required to discount alternative hypotheses such as PCR duplication of sequencing reads, reference-mapping bias, and copy number variants (CNVs), especially in immortalized cell lines that carry copy number aberrations^9,12^.

Although many non-motif-based features modulate TF binding affinity^8,13,14^, TF binding motifs provide insight into molecular mechanism when an ASB maps to a well-defined motif for the experimentally-targeted TF, and when its lower affinity allele disrupts this motif. Altered binding can disrupt a motif that is represented in publicly available databases such as JASPAR^15^, or a previously unknown TF binding motif predicted *de novo* directly from the ChIP-Seq data^16^.

To predict ASBs accurately, methods should call variants that meet four criteria: (i) they match known genotypes^9,11,12^, (ii) they meet stringent quality control and allele-dependent alignment of ChIP-Seq reads^17^, (iii) they lie within a ChIP-Seq peak, and (iv) they disrupt (or strengthen) a TF’s binding motif when it reduces (or increases) its DNA-binding affinity. Previous ASB prediction methods meet some but not all four criteria^9,10,18^. Furthermore, methods have not previously included a *de novo* motif discovery step, which limits the set of biologically-informative motifs to be investigated^17^. Prioritized ASBs that affect TF binding motifs can then facilitate genome-scale complex trait studies whose multiple testing burden is reduced due to the smaller number of prioritized variants.

Here, we describe *baal-nf*, a new computational framework that infers ASBs by processing and quality controlling ChIP-Seq data from a large number of studies that used genotyped cell lines, while accounting for known biases. Mapping these ASBs to known and *de novo* motifs then permits *baal-nf* to infer the TF mechanism by which binding is disrupted at these loci. To achieve our goals, we use nextflow to build an end-to-end pipeline, integrating tools for read and alignment quality control, alignment of high-quality reads, *de novo* motif discovery using NoPeak^16^ and inference of ASBs using BaalChIP^9^. BaalChIP is a fully-Bayesian approach that infers ASBs from ChIP-Seq data, and has been shown to accurately correct for biases due to copy number aberrations present in cancer cell lines.

We showcase *baal-nf* by its prediction of 298,783 ASBs from 558 TFs, and map these to 374 known and *de novo* motifs using publicly available ENCODE and ChIP-ATLAS data from 46 genotyped human cell lines. Among these are 1,960 high-quality, mechanistically *w*ell-understood ASBs. These are sites with strong evidence of binding, one of whose alleles both lowers a TF’s DNA-binding affinity and disrupts its binding motif. Consistent with their functionality, high-quality ASBs are more evolutionarily conserved across species, and are enriched for known trait associations and molecular quantitative trait loci (molQTL), supporting their prioritization in studies seeking causal trait-altering DNA variants.

## Results

### *baal-nf* workflow

Our aim was to create a workflow that infers ASBs, while correcting for sources of biological and technical bias, and that assigns their confidence based on disruption of known or *de novo* inferred TF motifs. To apply this tool across a large set of studies, the workflow needed to be scalable, parallelizable and reproducible, with adaptability to new computational settings. To achieve this, we used nextflow, a workflow management system that allows implementation of complex data analysis workflows and that can be run easily across various computational platforms. Docker containers were implemented to obtain reproducible software environments at each step in the workflow. The *baal-nf* workflow implements and builds upon BaalChIP, a Bayesian statistical approach that calls ASBs in cancer genomes^9^, as detailed below.

*baal-nf* requires three sets of input data: (1) ChIP-Seq FASTQ data from genotyped cell lines, (2) reference allele frequencies (RAFs) for all heterozygous SNPs in each cell line-of-interest, and (3) BED (Browser Extensible Data) files containing the ChIP-Seq peak calls for a given sequencing run [Figure 1A]. FASTQ files are first quality controlled by filtering out reads with low sequencing quality or reads harboring repetitive DNA, such as rDNA^19,20^ [Figure 1B, steps 1-2]. Remaining reads are then aligned to the human reference genome and duplicate reads marked as potential PCR duplications, before inferring ASBs with BaalChIP^9^ [Figure 1B, steps 3-5]. BaalChIP estimates the Corrected Allelic Ratio (CAR), the preference of TF binding to either the reference or alternate allele at heterozygous sites after correcting for biological and technical biases [Figure 1A] (see Methods). A CAR estimate of 0.5 indicates no preference for TF binding to either allele; an estimate between 0.5 and 1 indicates higher TF affinity for the reference (REF) allele; and an estimate between 0 and 0.5 indicates higher affinity for the alternate (ALT) allele.

**Figure 1:**
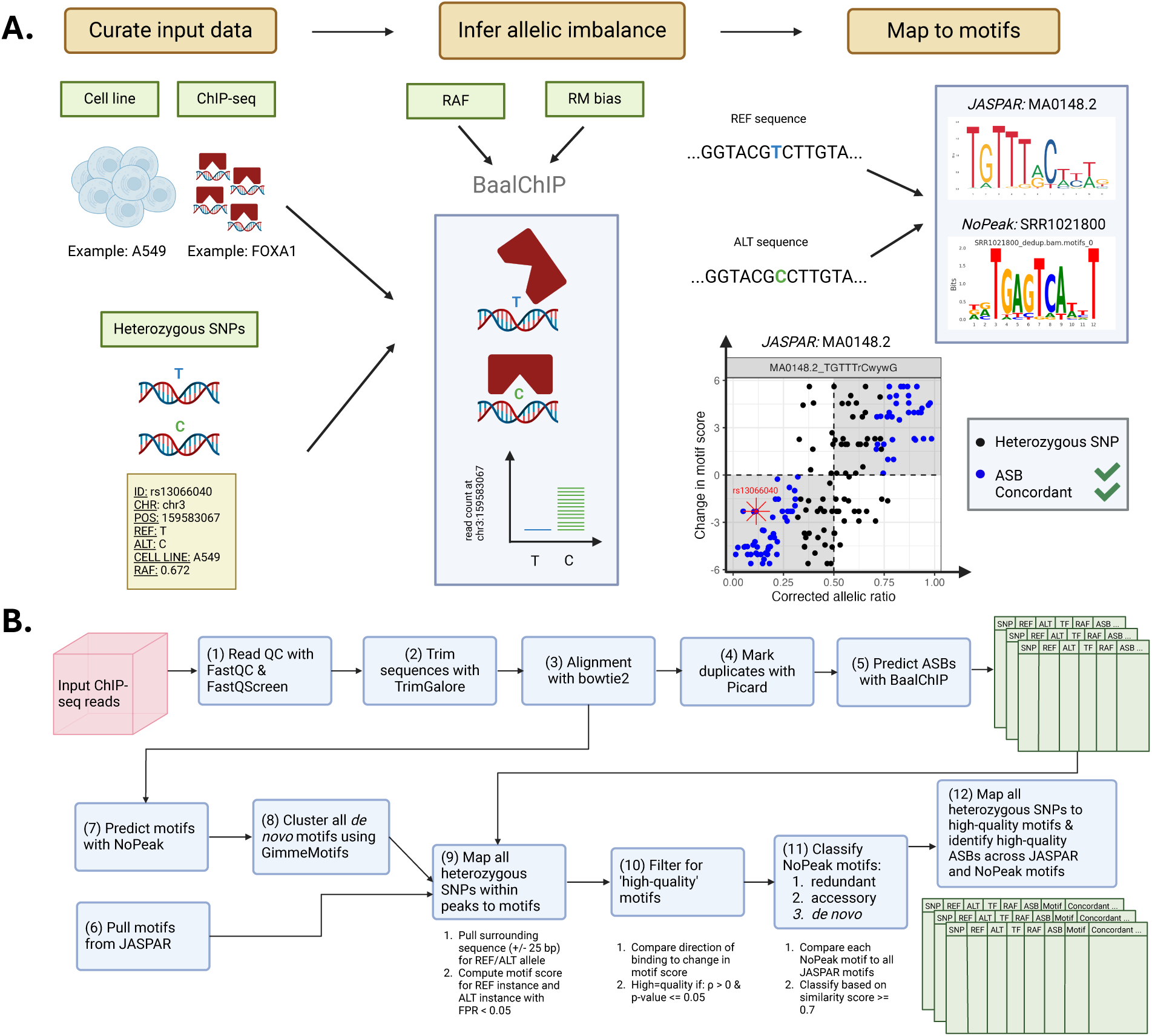
*baal-nf* workflow. **(A)** Workflow for identifying high-quality ASBs through *baal-nf*. ChIP-sequencing data from genotyped cell lines is used to infer ASBs at heterozygous SNPs using BaalChIP, a beta-binomial model that takes into account biological bias due to copy number aberrations (through the reference allele frequency (RAF)) and technical aspects such as reference-mapping (RM) bias. Identified ASBs are further characterized by mapping them to JASPAR and NoPeak motifs for a given TF. High-quality ASBs show concordance in the corrected allelic ratio (CAR) and motif score difference (MSD) as shown by the blue points in the bottom-right scatter plot. Data points are coloured blue when a heterozygous SNP is (1) an ASB, and either (2a) CAR > 0.5 and MSD > 0 (i.e., REF allele strengthens binding), or (2b) CAR < 0.5 and MSD < 0 (i.e., ALT allele strengthens binding). **(B)** Bioinformatic workflow for processing and quality-controlling data, acquiring and identifying motifs, and how heterozygous SNPs are mapped to these motifs and further classified.

Inferred ASBs are mapped to two types of TF motifs: known motifs for the experimentally-relevant TF (e.g., MA0148.2 for FOXA1), and *de novo* motifs that are enriched in the ChIP-Seq read data (e.g., SRR1021800) [Figure 1A]. Known motifs for the relevant TF are downloaded from the JASPAR database^15^ for use during the mapping procedure [Figure 1B, step 6]. *De novo* motifs are called from ChIP-Seq reads using NoPeak, a k-mer-based motif discovery method that predicts motifs from global read distribution profiles, without requiring background ChIP-Sequencing samples for motif discovery^16^ [Figure 1B, step 7] (Methods). NoPeak motifs are further clustered, combining information across highly-similar motifs and removing redundancy within the NoPeak motif set.

Only heterozygous SNPs that lie within ChIP-Seq peaks are considered at this stage, due to their strong evidence for TF binding, while we seek to derive a high-quality motif set. This set of heterozygous SNPs are mapped to all JASPAR and NoPeak motifs, resulting in motif scores for each of the REF and ALT alleles [Figure 1B, step 9] (see Methods). The difference in scores (the Motif Score Difference, MSD) indicates whether the REF or ALT allele better matches the motif-of-interest.

Motifs that best explain allelic imbalance of TF binding are defined as “high-quality motifs”: those in which mapped alleles with higher binding affinity (inferred from ChIP-Seq reads) show a significant tendency to be associated with stronger motif instance (i.e., with more positive MSD). More precisely, to be “high-quality” a motif has a significant and positive correlation (Spearman’s correlation coefficient, SCC) between the CAR derived from BaalChIP and the calculated MSD, considering only heterozygous SNPs in peaks whose surrounding sequence match this motif well [Figure 1B, step 10] (see Methods).

To characterize NoPeak motifs, we compare them against all known JASPAR motifs, defining a match between them based on motif similarity metrics. Motif similarity is assessed using the approach outlined in Grau et al.^21^ and Kielbasa et al.^22^ [Figure 1B, step 11] (Methods). Using this approach, high-quality NoPeak motifs are classified into one of three groups: (1) redundant motifs – those that are good matches to known motifs in the JASPAR database for the targeted TF, (2) accessory motifs – those that are good matches to known motifs in the JASPAR database that, however, are not linked to the ChIPped TF, and (3) *de novo* motifs – those that are poor matches to JASPAR motifs. After identifying NoPeak motifs similar to, and thus redundant for, JASPAR motifs, these are discarded from further analysis.

Once the set of high-quality motifs is derived, *baal-nf* maps the entire set of heterozygous SNPs back onto these motifs [Figure 1B, step 12]. The workflow first identifies “high quality ASBs”: those whose altered TF binding within a ChIP-Seq peak can be explained mechanistically by concordant disruption to a relevant, accessory or *de novo* TF binding motif. More specifically, these ASBs map to sequences matching a high-quality motif (for REF and ALT alleles) and that, additionally, show concordance between their direction of binding affinity (i.e., CAR) and their change in motif score (i.e., MSD) [Figure 1A; highlighted as blue data points]. Next, *baal-nf* identifies “low quality ASBs”: those that meet each of these criteria, except that they are located outside of a ChIP-Seq peak. We did not wish to discard all ASBs outside of peaks because peak-calling algorithms vary in performance across different binding mechanisms^23^. All remaining ASBs not meeting these criteria or that did not map to a high-quality motif were defined as “unclassified ASBs”. Further below, we justify naming ASBs as “high quality” based on evolutionary information and associations to quantitative trait loci (QTLs).

### Redundant, accessory and *de novo* motifs found for FOXA1

To exemplify these steps, we next describe how we applied *baal-nf* to a single TF, namely FOXA1.

FOXA1 belongs to the FOXA subfamily of winged helix transcription factors that bind and open chromatin, thereby facilitating access for other transcription factors^24^. FOXA1 also binds cooperatively to other nuclear receptors^25,26^ to enable transcriptional activation, and is associated with epigenetic regulation via DNA demethylation^27,28^.

FOXA1 binds DNA with high specificity, with binding affinity being highly dependent on minor variation of nucleotides within binding sites^29^.

To predict FOXA1 ASBs we used *baal-nf* and 156 ChIP-sequencing samples, 6 genotyped human cell lines and 270,587 heterozygous SNPs. Of 422,090 unique SNP-cell line pairs, 15,823 (3.7%) were predicted by BaalChIP to show allelic bias, with 420 of these replicated across two or more cell lines. Further, 9,214 (2.2%) lay within ChIP-Seq peaks whose sequences were good matches to JASPAR and/or NoPeak motifs for FOXA1.

All 4 JASPAR motifs for FOXA1 were high-quality motifs (i.e., showed concordant change in motif score and allelic ratio, [Figure 2A]) and 11 of 135 NoPeak motifs were high-quality. High-quality JASPAR or NoPeak motifs tended to have higher information content than low-quality motifs [Figure S1]. Of the 11 high-quality NoPeak motifs, 1 was redundant, 2 were accessory motifs, and 8 were *de novo*. The redundant NoPeak motif (Average_150) showed a higher similarity score (0.89) to the FOXA1 JASPAR motif MA0148.4 than other JASPAR motifs [vertical dashed red line, Figure 2B].

**Figure 2:**
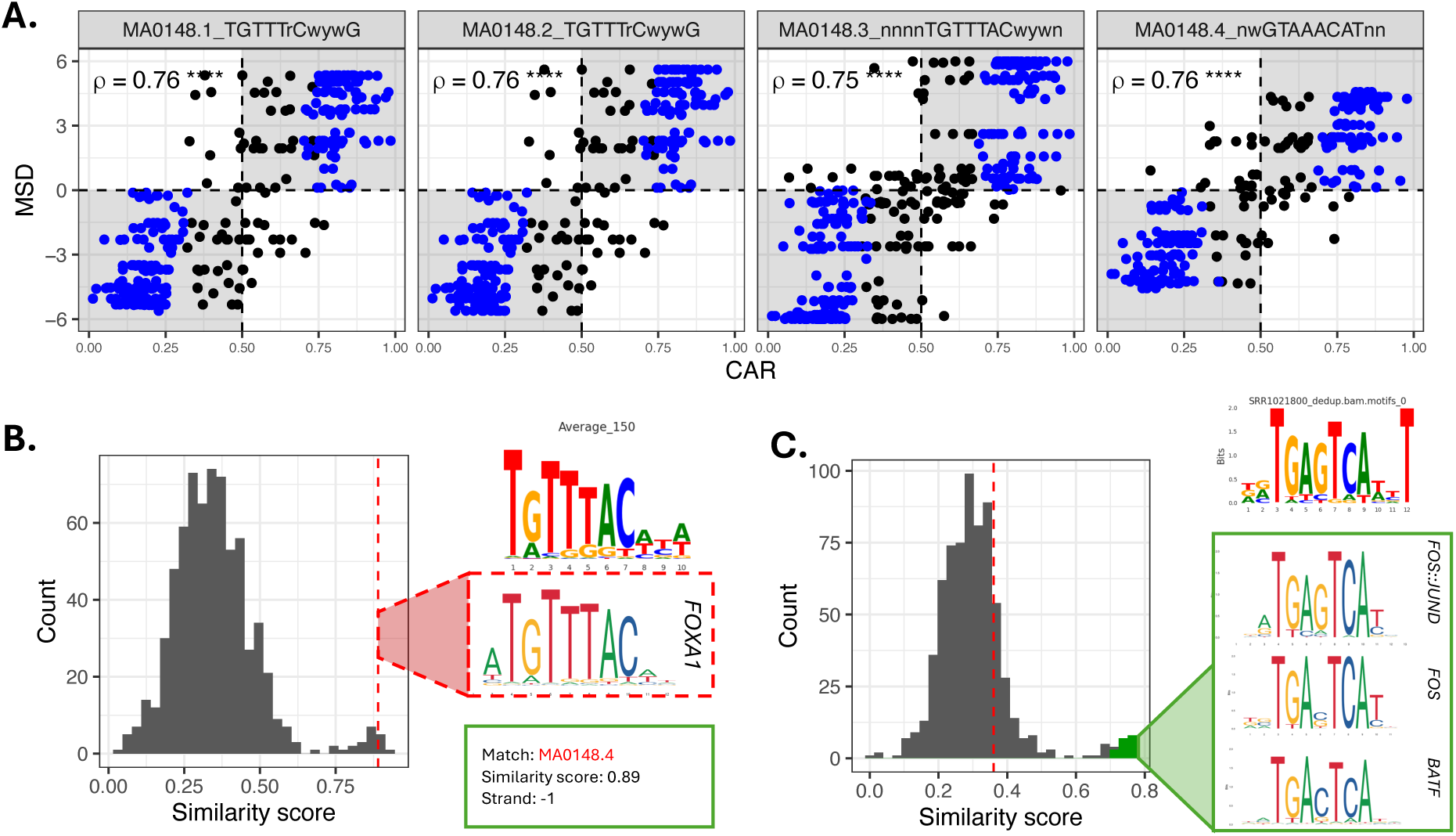
FOXA1 allele-specific binding inferred using *baal-nf*. **(A)** All four JASPAR motifs for FOXA1 are high-quality, showing a significant and positive SCC (ρ) between the CAR (x-axis) and MSD (y-axis). ρ is labeled on the plot, with significance levels **** representing a p-value < 2×10^-16^. Each scatter plot represents a unique JASPAR motif for FOXA1. **(B)** Redundant NoPeak motif identified for FOXA1 through *baal-nf*, where the NoPeak motif (“Average_150”) yielded a high similarity score of 0.89 (vertical dashed red line) to MA0148.4, a FOXA1 JASPAR motif, when compared to similarity scores for all JASPAR motifs, shown here as a histogram. **(C)** Accessory NoPeak motifs identified in FOXA1 ChIP-sequencing data, where the identified NoPeak motif (SRR1021800_motifs.0) yielded a high similarity score to FOS/JUN/BATF motifs (green cluster), and a low similarity score to FOXA1 JASPAR motifs (shown by dotted red line).

One of the 2 accessory motifs (SRR1021800_motif_0) is a good match to FOS/JUN heterodimer and BATF motifs [Figure 2C, Table S1], but not to FOXA1 motifs. FOS, JUN and BATF are all members of the AP-1 family of transcriptional activators, known to form heterodimers with one another and to be in TF-coordinated complexes with FOXA proteins^30^. FOS has also been proposed to activate FOXA1 through the ERBB2 signaling pathway^31^. The other accessory motif (ENCFF774LQB_motif_0) is a good match to FOXP2, another member of the FOX family of TFs. These results demonstrate *baal-nf*’s ability to discover mechanistically-predictive TF binding motifs.

A total of 1,246 ASB events (7.9%) mapped to the 14 high-quality, non-redundant motifs (10 NoPeak, 4 JASPAR), with 224 being high-quality ASBs and 633 being low-quality ASBs. Across these 14 motifs, a median of 60 low-quality ASBs and 15 high-quality ASBs were identified [Table S2]. High-quality ASBs were discovered across JASPAR and NoPeak motifs in approximately equal measure, with 58% (129/224) mapped to JASPAR motifs, and 42% mapped to NoPeak motifs (16 to accessory motifs and 79 to *de novo* motifs) [Table S3, Figure S2, Figure S3]. This approximate doubling of high-quality ASB predictions for FOXA1 highlights the importance of including *de novo* motif discovery in this workflow. High-quality ASBs for FOXA1 were associated, on average, with 2.7 human physiological traits at a p-value threshold < 5 × 10^-8^ and 1.8 expression QTLs (eQTLs) [Figure S4] (Methods).

### *baal-nf* predicts 1,960 high-quality binding variants across 558 TFs

We applied *baal-nf* to 6,017 ChIP-Seq data sets for 558 TFs and 46 genotyped cell lines (Table S4), making up 1,056 TF-cell line groups. This resulted in 298,783 inferred ASBs, with 8,968 (3.00%) mapping to high-quality TF motifs. Of these, the binding change affinity for 5,063 (1.69%) was concordant with disruption to a relevant motif, and 1,960 (0.66%) passed the criteria (defined above) necessary to be assigned as high-quality [Figure 3A, Table S5, Table S6]. Among 3,014 JASPAR and NoPeak motifs investigated, 374 were high-quality [Figure S5]. Compared to low-quality motifs, high-quality motifs yielded consistently larger, positive Spearman’s correlation coefficients between CAR and MSD, indicating that they are more explicable of binding variation for investigated SNPs [Figure S6].

**Figure 3:**
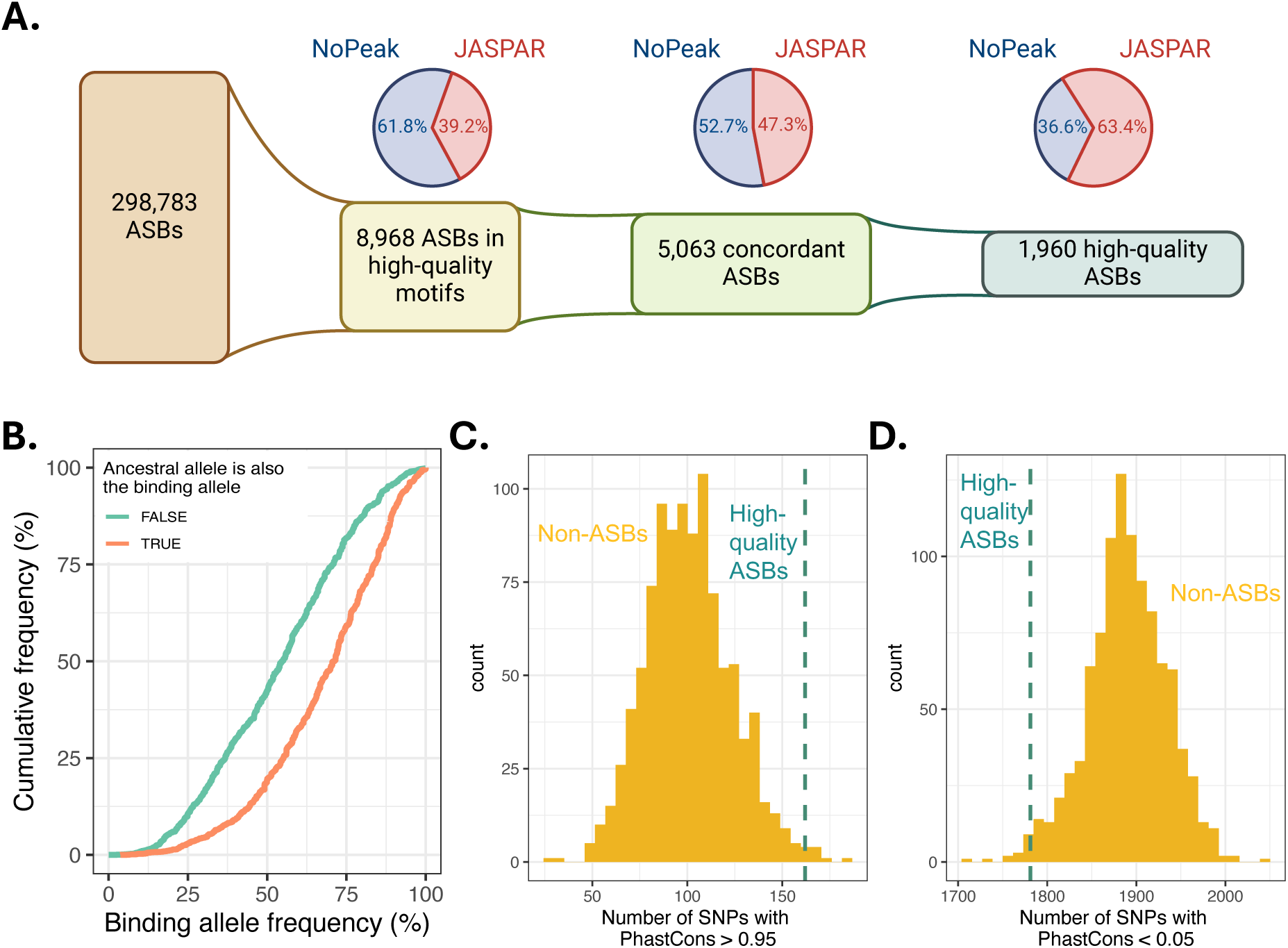
ASB set derived across 558 TFs and 46 genotyped cell lines, resulting in a subset of high-quality ASBs. **(A)** From the complete set of ASBs predicted by BaalChIP, 3% of these map within high-quality motifs, 5,063 display concordant behaviour in direction of inferred binding from BaalChIP (CAR) and disruption to a relevant motif (MSD), and 1,960 are deemed to be high-quality. **(B)** Empirical cumulative density function (ECDF) of frequencies of the binding allele in the non-Finnish European population from gnoMAD, split by whether the binding allele is the ancestral or derived allele. A Wilcoxon rank sum test comparing the means between the groups (FALSE and TRUE) show a significantly higher binding allele frequency when the ancestral allele is also the binding allele (TRUE), with a p-value of 5.4 × 10^-66^. **(C)** High-quality ASB sites (turquoise dashed line) when compared to non-ASB sets (yellow histogram) are significantly better conserved, as indicated by a higher number of SNPs with PhastCons scores > 0.95. **(D)** We find the opposite effect for non-conserved bases, with a significantly lower number of non-conserved bases present in the high-quality ASB set, represented by a PhastCons scores < 0.05.

High-quality ASBs (n=1,960) were predicted by *baal-nf* for 86 out of 558 TFs in 46 genotyped cell lines. JASPAR motifs contributed most to this high-quality subset (average of 22 high-quality ASBs per TF), followed by NoPeak *de novo* motifs (average of 18 high-quality ASBs per TF) and NoPeak accessory motifs (average of 13 high-quality ASBs per TF) [Figure S7]. As with the FOXA1 example (above), inclusion of NoPeak motifs substantially expanded the set of high-quality ASBs, as a result of both accessory and *de novo* motifs being discovered across TFs [Figure S8].

Variation in TF-binding affinity may not alter downstream molecular, cellular or organismal traits, and thus may not be functional^32^. To assess functionality of the high-quality ASB subset, we considered whether they had been subject to evolutionary selection, have known associations with human traits or are QTLs for transcript abundance or splicing (i.e., eQTL and sQTL, respectively).

For the 1,960 high-quality ASBs, we hypothesized that the allele associated with stronger TF binding affinity, i.e., the “binding allele”, would more often be the ancestral allele that has been retained in the population at higher frequency than the lower affinity, derived allele. Indeed, we discovered that when the binding allele is the derived allele then its population frequency is typically lower than when the binding allele is the ancestral allele (p-value=5.4 × 10^-66^) [Figure 3B]. This is consistent with positive selection of the binding allele and/or negative selection of the lower affinity allele.

Next, we showed that high-quality ASB sites tend to be evolutionarily well conserved. For this analysis, we generated 1000 comparator sets of “non-ASBs”, defined as randomly sampled high read coverage heterozygous SNPs that, when tested by *baal-nf*, failed to show significant allelic bias in any cell line, and for any TF (Methods) [Table S7]. We define high read coverage here as ≥ 100 reads mapping to the region containing the SNP. SNPs in these non-ASB sets were matched on minor allele population frequencies to those of high-quality ASBs, and had the same number of SNPs as the high-quality set (Methods).

SNP positions in both groups were scored for evolutionary conservation across 30 eutherian mammals using PhastCons^33^. PhastCons scores range between zero – no conservation of that SNP across the species-of-interest – and one, indicating complete conservation. The number of highly conserved bases (PhastCons score > 0.95) was significantly larger in the high-quality ASB set (empirical p-value = 8.0 × 10^-3^), and the number of non-conserved bases (PhastCons score < 0.05) was significantly lower in the high-quality ASB set (empirical p-value = 1.3 × 10^-2^) [Figure 3C-D].

Compared with non-ASBs, we found that high-quality ASBs also had: (i) a significantly higher number of known variant-human trait relationships (p-value = 2.75 × 10^-3^), (ii) more eQTLs (p-value = 3.0 × 10^-9^), and (iii) a higher maximum OpenTargets V2G (Variant-to-gene) score when associating each variant with a downstream gene (p-value = 1.9 × 10^-9^), but no significant difference in the number of sQTLs (p-value = 0.97)^34^. As expected, minor allele frequencies were not significantly different between our non-ASB set and high-quality set, mitigating ASB discovery biases [“count100” in Figure S9A].

Findings were also robust to different count thresholds used to define high coverage SNPs for the random samples of non-ASBs [Figure S9B-E, Figure S10]. Both evolutionary conservation analyses and trait/QTL-association analyses were consistent when using a more lenient minimum read coverage threshold of 50 to derive our 1000 non-ASB comparator sets (results for “count50” are shown in Figures S9B-D and S10).

With the same analyses, “low-quality ASBs” were found not to be more evolutionarily conserved compared to their non-ASB sets [Figure S11]. They were, however, significantly associated (p-value = 2.0 × 10^-3^) with a higher number of human traits at a p-value threshold < 5 × 10^-8^, but not for the numbers of colocalized eQTLs or sQTLs, or the maximum V2G score [Figure S12, Table S8, Table S9].

In summary, by applying *baal-nf* across a large set of TFs, we have generated a new database of 298,783 ASBs, and have identified a subset of 1,960 high-quality ASBs. For these high-quality ASBs, the differential binding mechanism, inferred using high-quality motifs, provides high confidence for functionally-significant altered binding at these loci. When compared to non-ASB SNPs, this high-quality ASB set exhibited greater conservation across species, a higher number of known associations with traits, and greater colocalization with molecular QTLs.

## Discussion

Identification of trait-causal variants would yield insights into fundamental biology that could aid development of therapeutic interventions. Nevertheless, such variants are challenging to identify due to linkage disequilibrium, whereby neighboring SNPs are often co-inherited, leading to significant variant-trait associations that mainly reflecting correlation, not causation. Assigning molecular mechanisms to SNPs is critical for prioritization of variants that are truly causal^35,36,37,38^. Extensive cataloguing of molecular QTLs is required to refine the evidence for how finely-mapped SNPs can mechanistically explain trait variation.

Substantial efforts have been made to identify and annotate cis- and trans-expression QTLs and splicing QTLs through databases like eQTLGen^39^, GTEx^40^, eQTL Catalogue^41^, single-cell eQTLGen Consortium^42^, as well as integrated platforms such as QTLbase2^43^ and OpenTargets Genetics^34,37^. Less effort has been expended on TF binding QTLs: such resources are scarce and are seldom integrated into these platforms. Databases for ASBs like AlleleDB^44^ and ADASTRA^10^, with extensions such as ANANASTRA^45^, have tried to address this gap. Here, we sought to help narrow the information gap and provide a tool for large-scale identification of new and high-quality ASB datasets.

Nevertheless, the ASBs reported in this study are far from being complete. This is because *baal-nf* can only infer ASBs for heterozygous SNPs in genotyped cell lines with ChIP-Seq data. ASB incompleteness is evident from the near disjoint sets of ASBs inferred by *baal-nf* (n=298,783 ASBs) and by ADASTRA (n=264,318 ASBs)^10^ [Figure S13A] caused by minimal overlap in the TFs and cell lines investigated by the two methods, and by ADASTRA not requiring ASBs to have a priori known genotypes [Figure S13B-C]. This shortfall can be addressed by investigators applying *baal-nf* to new ChIP-Seq datasets. Even with application to new datasets, low coverage of ChIP-Seq reads at heterozygous sites will impede our ability to call ASBs, as there may be insufficient power to do so.

Prediction incompleteness is also evident after comparing *baal-nf* ASBs to those predicted from SNP evaluation by Systematic Evolution of Ligands by EXponential enrichment (SNP-SELEX) data in GVATdb^46^. Compared with GVATdb, *baal-nf* tested an order of magnitude higher number of TF-SNP pairs (17.5-fold higher) and called an order of magnitude higher numbers of ASBs (13.5-fold), demonstrating similar ASB call rates across studies. A very small percentage of TF-SNP pairs were tested in both strategies (with the same REF/ALT alleles) - 3,170 of 1,374,477 (0.2% of GVATdb) and 3,170 of 23,997,518 (0.01% of *baal-nf*).

Within this set of 3,170 non-redundant TF-SNP pairs, there were 4,435 ASB tests in total due to multiple experiments and/or cell lines. Of these, 3,083 (97.3%) were called as non-ASBs in both; 83 (2.6%) were discordant, being called as a non-ASB by *baal-nf* and as an ASB in GVATdb, or vice versa; and 4 (0.1%) were called as an ASB by both *baal-nf* and in GVATdb. Note that the same SNP/TF pair could be called as both a non-ASB and ASB in both databases due to variation across experiments, biological contexts (i.e., different cell lines) and statistical coverage (i.e., read depth). The low numbers of TF-SNP pairs investigated in both these studies demonstrates how far genomics research has yet to progress before testing anywhere near the full complement of possible TF binding QTLs in the human genome.

We note that *baal-nf* can easily be extended to predict allelic imbalance from any other sequencing-based method, including CUT&RUN and CUT&TAG, or RNA-protein based approaches such as Cross-linking and immunoprecipitation followed by sequencing (CLIP-Seq).

*baal-nf* is an open-source software, available on GitHub, which can be used to derive new ASB sets and high-quality ASBs from ChIP-Seq datasets. It supports end-to-end processing of raw sequencing data, inference of ASBs and characterization of these ASBs with respect to binding mechanism. For large-scale application, a high-performance compute (HPC) cluster is recommended for efficient parallelization of the workflow. A major challenge in genomic research is the ability to make research portable and reproducible^47^. To achieve this, we rely on nextflow and singularity, allowing large-scale implementation of this tool with stable and reproducible software environments^48^. Limitations of *baal-nf* include BaalChIP’s reliance on the hg19 genome assembly, and its current dependence on ChIP-sequencing data. Furthermore, non-JASPAR reference motifs are not currently integrated into the workflow. This tool can be extended in the future for compatibility with additional genome assemblies, motif databases, and other sequence-based methods that measure allelic imbalance.

We hope that others will add to this initial discovery set of high-quality ASBs by applying *baal-nf* to further data sets. All such high-quality ASBs will be useful when prioritizing non-coding regulatory variants as causal of complex trait variation and disease risk.

## Conclusions

Of the 298,783 ASBs ASBs reported in this study, 0.66% (n=1,960) of these were “high-quality”. We proposed this high-quality set as strong candidates for causal trait-altering DNA variants, as demonstrated by enrichment for known variant-trait relationships, colocalization with gene expression QTLs and strong evolutionary conservation.

Although limited in size, this high-quality set’s variants are mechanistically well-understood and have evidence for altered binding across multiple datasets. This is because the method requires: (1) strong evidence for the called genotype at heterozygous SNPs, (2) strong evidence for allelic imbalance, (3) orthogonal evidence for altered binding by disruption of a TF binding motif, and (4) strong evidence for binding by residing within a ChIP-Seq peak. By proposing a high-quality set, we enable researchers to prioritize these variants in large-scale genomic studies.

## Methods

### Read QC, alignment and inferring ASBs

For *baal-nf*, read quality is assessed using FastQC (version v0.11.9) and FastQScreen (version v0.14.0) and then trimmed using TrimGalore! (version v0.6.7), filtering out any reads lying below a minimum quality threshold (default parameters in TrimGalore!)^19,20^. High-quality reads are aligned using bowtie2^49^ (version v2.3.5.1) to hg19, the human reference genome used in BaalChIP, duplicate reads are marked with Picard (version v2.23)^50^, and aligned BAM files are used to infer ASBs using BaalChIP^9^ (custom version modified from v1.1.1; provided at https://github.com/BAAL-NF/BaalChIP). BaalChIP models allelic counts with a beta-binomial model, correcting for CNVs using the RAF as a prior, and correcting for Reference Mapping (RM) bias, which is estimated directly from the data. An ASB is called only if the highest posterior density interval for the fraction of reads mapping to the REF allele does not include a value of 0.5.

Heterozygous SNPs predicted to be ASBs will have a value of “True” set in the output column “isASB”. The Corrected Allelic Ratio (CAR) will also be reported for each SNP, which describes preference in binding to either the REF (CAR > 0.5) or ALT (CAR < 0.5) allele, after correcting for the RAF and RM bias.

### Predicting motifs with NoPeak

Aligned BAM files are converted to BED files using bedtools (version v2.29.2), and NoPeak^51^ is used to compute score profiles for each k-mer of length 8, the NoPeak default setting. Briefly, a score profile is computed by estimating the density of reads which include a given k-mer, and computing the distance from the beginning of that read to the k-mer of interest. Profiles that are consistent with TF binding have an increased frequency of reads over the k-mer sequence. All k-mer profiles that indicate TF binding are combined based on sequence similarity, resulting in predicted motifs per-ChIP-sequencing sample. Low k-mer motifs derived from fewer than 10 k-mers are excluded from subsequent steps as they are generally low-complexity and not indicative of true binding motifs.

### Identifying non-redundant NoPeak motifs

Predicted NoPeak motifs are defined per-ChIP-sequencing sample, leading to likely redundant motifs across multiple samples ChIPped for the same TF. To remove redundant motifs, all NoPeak motifs discovered for a given TF are clustered using GimmeMotifs (version v0.17.2)^52^, using de Bruijn sequences to quantify motif similarity (cluster_motifs() function: trim_edges = True, metric = “seqcor”, threshold = 0.7, combine = “mean”). A minimum threshold of 0.7 is chosen based on the analysis using the “seqcor” metric in Grau et al^21^. This scores each motif against one another, trimming low information content bases at the edge of the motif, and clustering motifs together that have a similarity score greater than 0.7. Each motif cluster is then averaged across the length of the motif to obtain a final, clustered motif. The resulting motifs from NoPeak are named after the ChIP-sequencing sample in which it was discovered or, if clustered, are named Average_n, where n is a number randomly-generated by GimmeMotifs.

### Mapping heterozygous SNPs to JASPAR and NoPeak motifs

Heterozygous SNPs are mapped to JASPAR and NoPeak motifs using GimmeMotifs (version v0.17.2)^52^. Sequence instances for the REF and ALT allele for each heterozygous SNP are derived by extracting +/- 25 bp around the position of the relevant SNP from the human reference genome assembly (here, hg19). Each instance is scored against a motif-of-interest (the motif score), which is represented as the log-odds of belonging to that motif compared to a random background sequence from the same reference genome, with a false positive rate less than 5 × 10^-2^. The Motif Score Difference (MSD) is computed by taking the difference between the REF motif score and the ALT motif score. An MSD of zero indicates that the REF allele does not disrupt the motif-of-interest compared to the ALT allele; a positive MSD indicates that the REF allele is more likely to belong to the motif-of-interest; and a negative MSD indicates that the ALT allele is more likely to belong to the motif-of-interest. “High-quality” motifs are determined by computing the Spearman’s Correlation Coefficient (SCC; rho) between the CAR and MSD across all heterozygous SNPs that map to a given motif, and then filtering for motifs with a significant, positive SCC (p-value < 0.05). Only such motifs are considered when identifying high-quality ASBs.

### Characterizing NoPeak motifs

To assess similarity of NoPeak motifs to known JASPAR motifs, we compute a similarity score for each NoPeak motif compared to all known JASPAR motifs (organism: Homo sapiens) using GimmeMotifs (version v0.17.2). We follow the approach outlined in Grau et al.^21^ and Kielbasa et al.^22^ using de Bruijn sequences to compute a similarity score.

Each step along the chosen de Bruijn sequence, representing every k-mer of length k (k=7, the default provided in GimmeMotifs), is scored against the first motif and second motif, resulting in two vectors of motif scores, termed score profiles. If the motifs are highly similar, then these score profiles are highly correlated, and have a strong, positive Pearson Correlation Coefficient (PCC). The PCC between score profiles is computed for every possible offset of the two motifs as well as the reverse complement, and the maximum score (termed “similarity score”) is taken. NoPeak motifs with a similarity score greater than 0.7 to a known JASPAR motif will either be classified as (1) redundant: the NoPeak motif matches the canonical JASPAR motif for the TF-of-interest, in which case it is removed from further analysis, or (2) accessory: the NoPeak motif matches a known JASPAR motif that is not the canonical binding motif Kielbasa et al^22^.

### ASB subsets – defining high and low-quality ASBs

Concordant ASBs are derived by filtering for all ASBs that map to a high-quality motif, and either (1) the CAR is > 0.5 and the MSD > 0 or (2) the CAR < 0.5 and the MSD < 0. This subset is further characterized by whether or not the SNP lies within a ChIP-Seq peak. ASBs that lie within ChIP-Seq peaks are termed “high-quality” and those that lie outside of ChIP-Seq peaks are called “low-quality”. As the same SNP can map to multiple motifs, we define an order in which to call SNPs concordant by first prioritizing those that lie within (i) JASPAR motifs, then (ii) NoPeak accessory motifs and then (iii) NoPeak *de novo* motifs.

### Deriving non-ASB sets

To compare sets of “non-ASBs” to the high-quality ASB set, all assessed heterozygous SNPs across all 46 genotyped cell lines were inspected, and any SNP (specifically rsID) for which an ASB was called by BaalChIP was removed from this set. This resulted in a final set of analysed SNPs for which an ASB was not called in any cell line assessed across all 558 TFs. Each heterozygous SNP in this set was then required to be “high coverage” (number of reads ≥ 100 at its site) to ensure sufficient coverage to confidently call this site as non-ASB. We also evaluated this workflow at minimum read threshold of 50 [“count50” in Figures S9-S11]. Every SNP in this non-ASB sample set was queried using the ENSEMBL REST API to pull the gnoMAD minor allele frequency for the non-Finnish European population. Non-ASBs’ rsIDs were randomly sampled with replacement from sets matched on minor allele frequency (MAF) within a 5% relative threshold, and this process was repeated across all high-quality SNPs to form a sample size of 2,400 non-ASB SNPs, a number matching the count of ASBs found in the high-quality set, including SNPs that were replicated across cell lines for the same TF. This sampling procedure was repeated 1000 times to generate 1000 non-ASB sets. A single “median” non-ASB set was also derived by selecting the SNP with the median MAF across the 1000 sampled datasets for each SNP in the total set of MAF-matched SNPs (n=2,400). This results in a single “median” non-ASB dataset matching the SNP set size of 2,400. This median set was used for querying statistics from OpenTargets, including the number of trait associations found at a p-value threshold of 5 × 10^-8^, the number of colocalized eQTLs and sQTLs, and the maxiumum V2G score for that variant. This same process was repeated for low-quality ASBs, with a set size of 3,473 to derive new non-ASB reference sets matched on MAF. See Figure S14 for a flow chart of this workflow.

### Evolutionary and functional genomics analysis of SNP groups

PhastCons scores for all SNPs assessed in 30-way bigwig files (for GRCh38) were pulled from UCSC^53^ and scores computed for SNPs in high-quality ASB and non-ASB groups using ENSEMBL’s variant effect predictor VEP (version 112)^54^. Variant-trait and variant-gene relationships were determined for each SNP group by querying OpenTargets Genetics^34,37^ using their GraphQL API. For each variant, every variant-trait and variant-gene relationship record was pulled, and the following values were computed for each variant: (1) number of traits that passed a significance threshold of p<5×10^-8^, (2) number of colocalized eQTLs, (3) number of colocalized sQTLs, and (4) maximum V2G score. Additional information was queried using the ENSEMBL REST API, including information about which allele was ancestral, as well as minor/major alleles in a population-of-interest and their respective frequencies in that population.

Here, we used the gnoMAD non-Finnish European population, and in cases where this data was not available, the 1000Genomes GBR population. See Figure S15 for a flow chart of this workflow.

## Declarations

### Ethics approval and consent to participate

Not applicable.

### Consent for publication

Not applicable.

### Availability of data and materials

All data generated or analysed during this study are included in this published article or are from publicly-available sources. All supplementary tables can be downloaded from the Open Science Framework (OSF) at https://osf.io/rwjec/ under the project osf.io/rwjec (v0.1.0; DOI 10.17605/OSF.IO/RWJEC). Accession IDs for FASTQ and BED files from ENCODE (at https://www.encodeproject.org/)^55^ & ChIP-ATLAS (at https://chip-atlas.org/)^56^ are described in Table S4. Additional ChIP-Seq FASTQ and BED files were used for the Vitamin D Receptor (VDR) from Gallone et al^57^. Results from this study are contained within Tables S5, S6, and S10. Table S5 contains all 298,783 predicted ASBs across 558 TFs, Table S6 contains all high-quality ASBs and queried information from ENSEMBL and OpenTargets. Table S10 contains all motifs explored in this study with associated metadata for each TF. All SNP-cell line mappings, including characterization with respect to high-quality motifs, are deposited on OSF under project osf.io/rwjec in the folder “Database”, separated by transcription factor. All code is deposited on GitHub at https://github.com/BAAL-NF/baal-nf and is publicly available. *baal-nf* version implemented in this manuscript is 0.9.0. All figures were generated using BioRender^58^. For the purpose of open access, the author has applied a creative commons attribution (CC BY) licence to any author accepted manuscript version arising.

### Competing interests

The authors declare that they have no competing interests.

### Funding

**Breeshey Roskams-Hieter** was supported by the Health Data Research UK & The Alan Turing Institute Wellcome PhD Programme in Health Data Science (Grant Ref: 218529/Z/19/Z). **Øyvind Almelid** was supported in part by an Alan Turing Institute award to Chris P Ponting (TU/ASG/R-SPEH-102). **Chris P Ponting** was supported in part by the Medical Research Council (MC_UU_00007/15).

### Authors’ contributions

**Breeshey Roskams-Hieter**: Conceptualization, Methodology, Software, Validation, Formal analysis, Writing – Original draft, Visualization, and Funding Acquisition. **Øyvind Almelid**: Conceptualization, Methodology, Software, Formal analysis, Data Curation, Writing – Review & Editing, and Supervision. **Chris P Ponting**: Conceptualization, Funding Acquisition, Methodology, Supervision, and Writing – Review & Editing.

## Acknowledgements

None to declare.

## Supplementary Figures

**Supplementary Figure S1:**
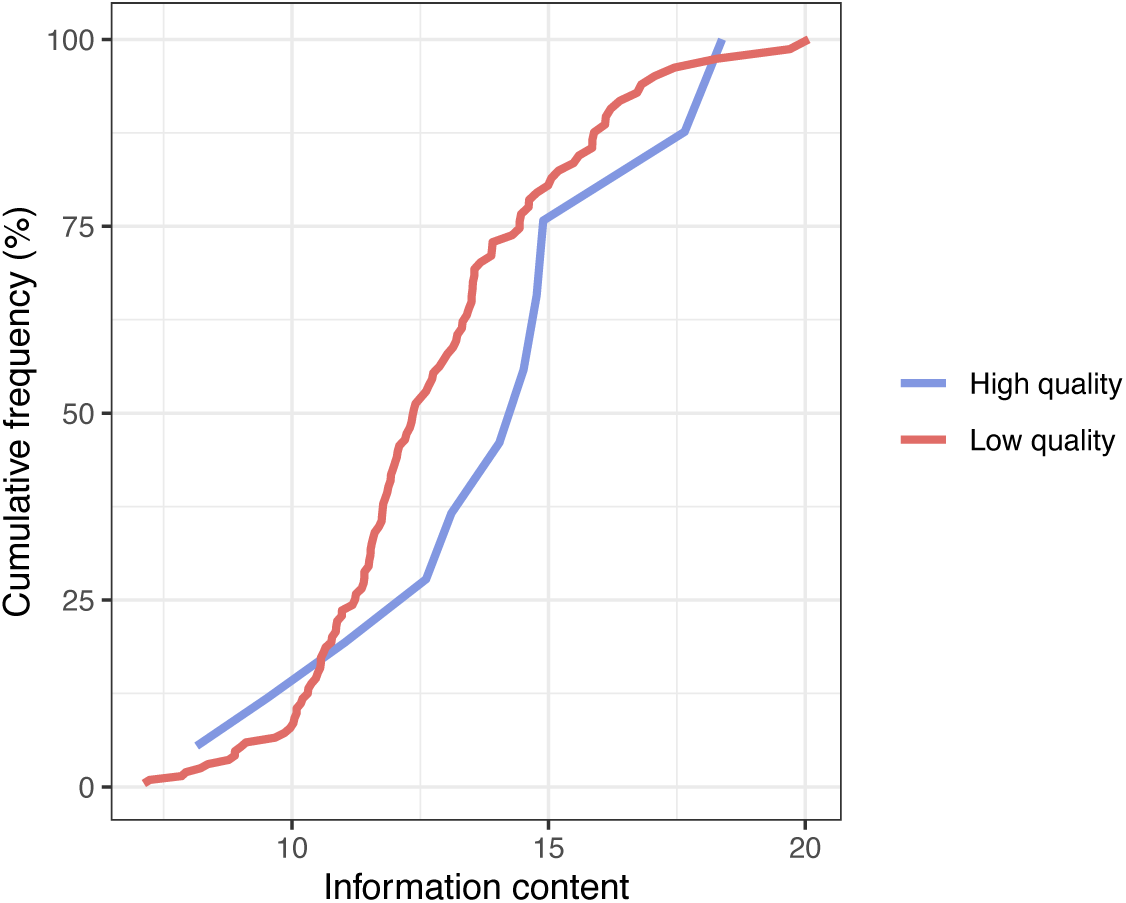
Properties of high-quality motifs for FOXA1. Empirical cumulative density frequency (ECDF) plot showing information content of high-(blue) versus low-(red) quality motifs found for FOXA1. Information content (x-axis) is measured in bits.

**Supplementary Figure S2:**
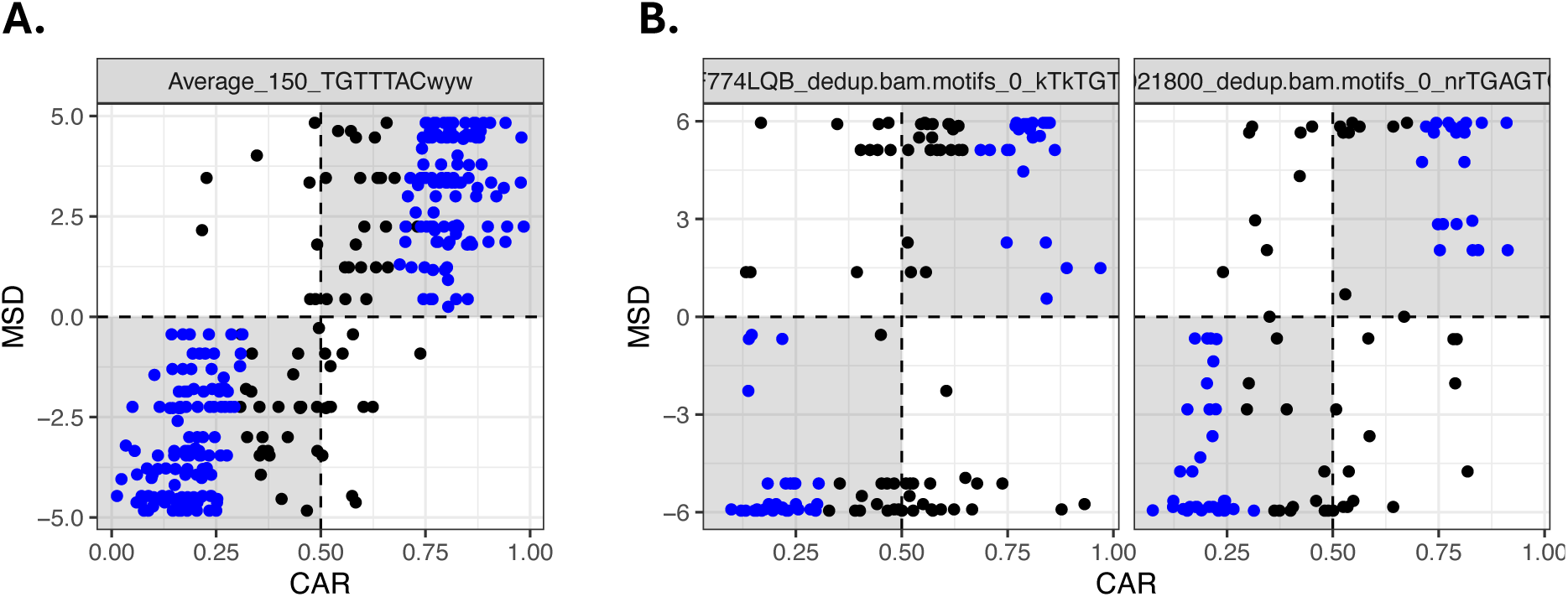
Redundant and accessory motifs discovered for FOXA1. Scatter plots for **(A)** redundant and **(B)** accessory motifs found for FOXA1 via NoPeak. High-quality ASBs are coloured in blue. CAR (x-axis) is the corrected allelic ratio calculated by BaalChIP for a heterozygous SNP lying within a ChIP-Seq peak; MSD (y-axis) is the motif score difference for each SNP mapped to the labeled motif.

**Supplementary Figure S3:**
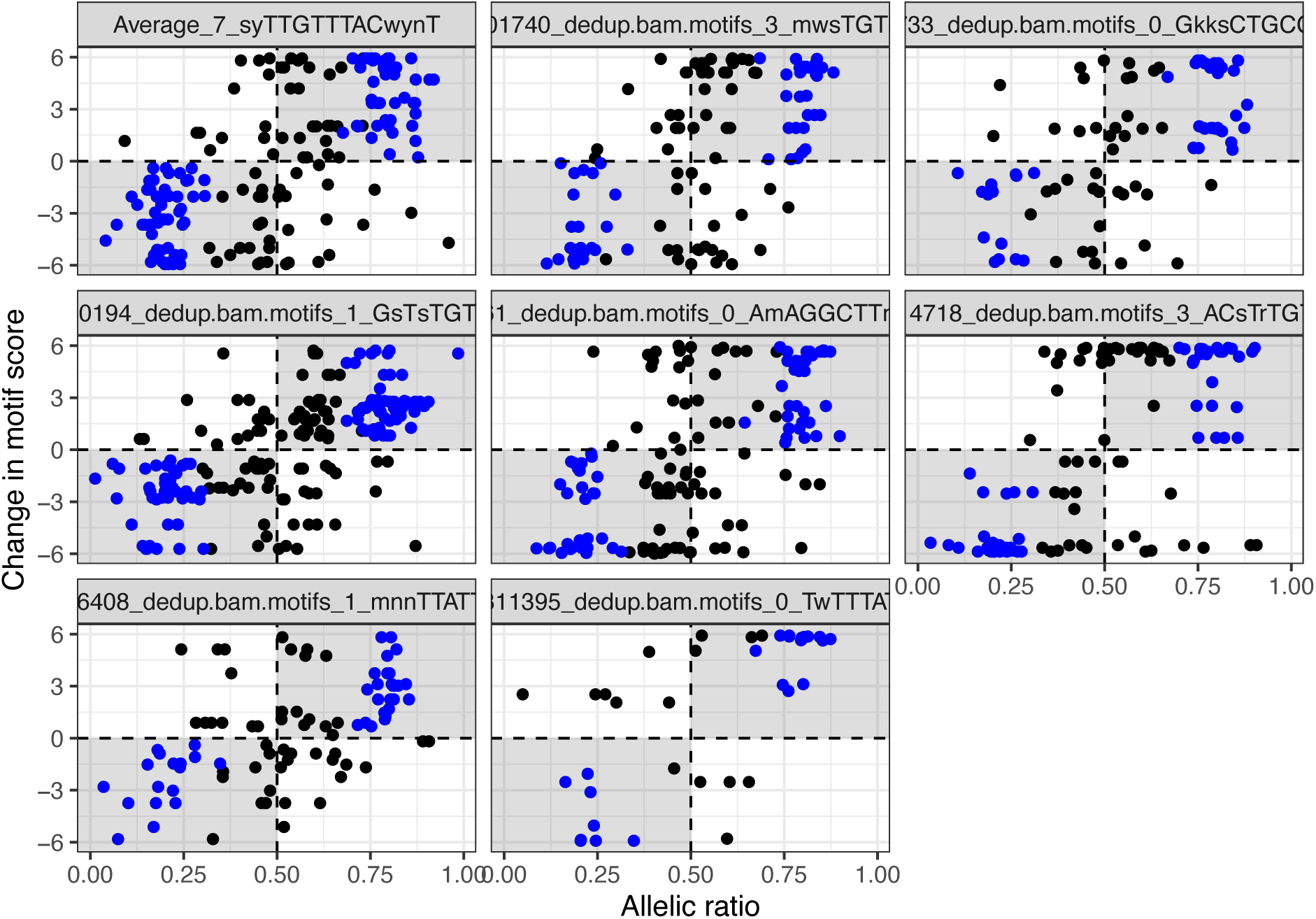
de novo motifs found for FOXA1 via NoPeak. High-quality ASBs are coloured in blue. CAR (x-axis) is the corrected allelic ratio calculated by BaalChIP for a heterozygous SNP lying within a ChIP-Seq peak; MSD (y-axis) is the motif score difference for each SNP mapped to the labeled motif.

**Supplementary Figure S4:**
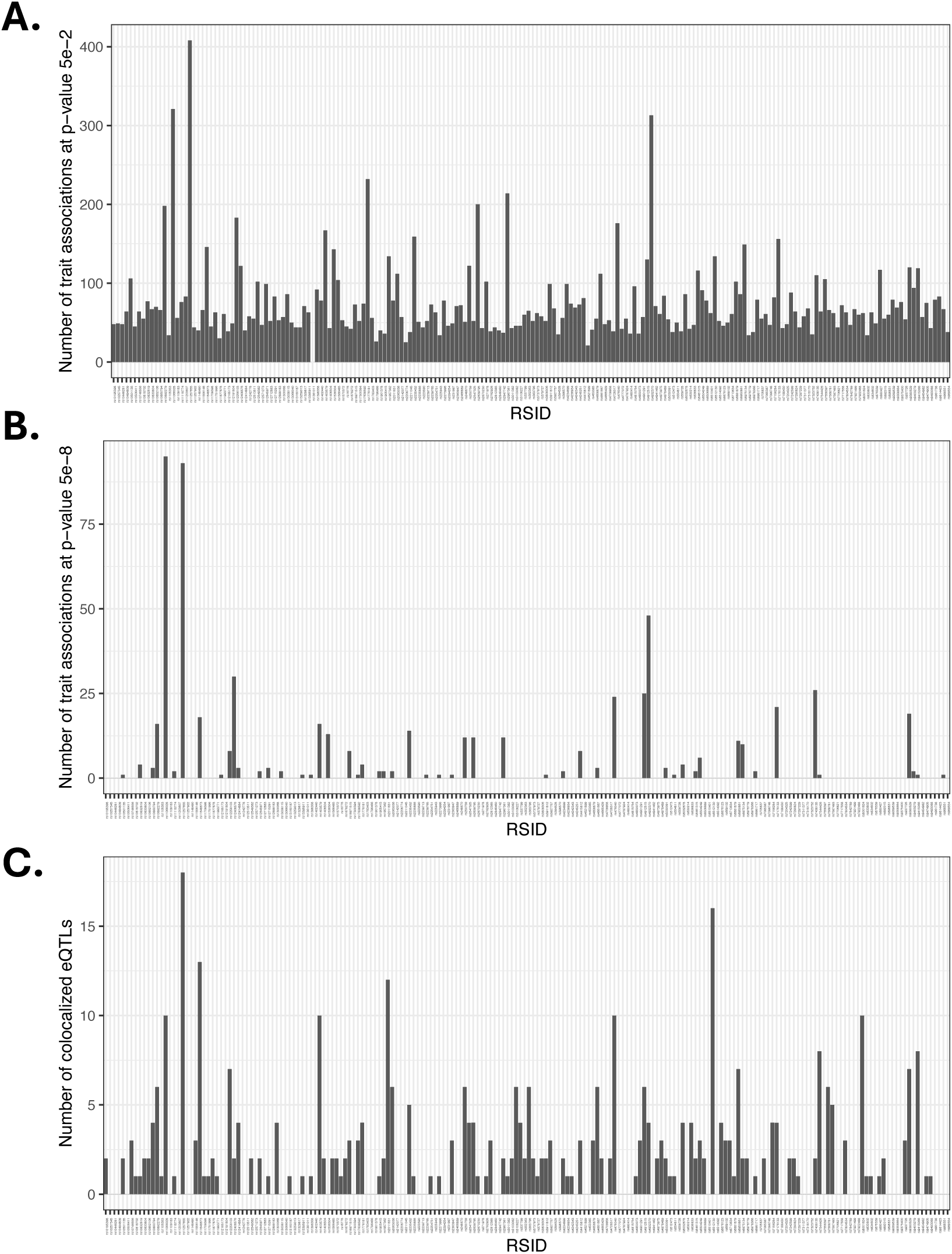
Number of associated traits & eQTLs for high-quality ASBs predicted for FOXA1. These values were obtained by querying for the variant in OpenTargets Genetics and summarizing statistics for each category – **(A)** Number of trait associations that passed a p-value threshold of 5 × 10^-2^, **(B)** Number of trait associations that passed a p-value threshold of 5 × 10^-8^, and **(C)** Number of colocalized eQTLs.

**Supplementary Figure S5:**
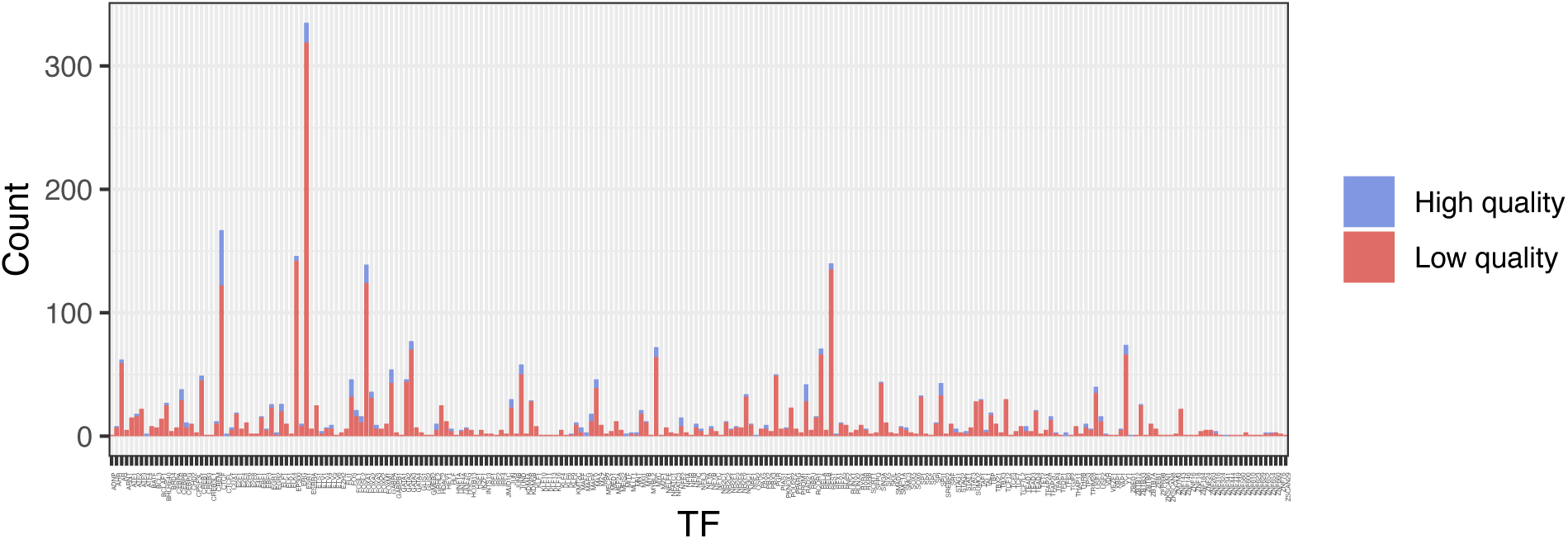
Number of high and low-quality motifs detected across 558 TFs.

**Supplementary Figure S6:**
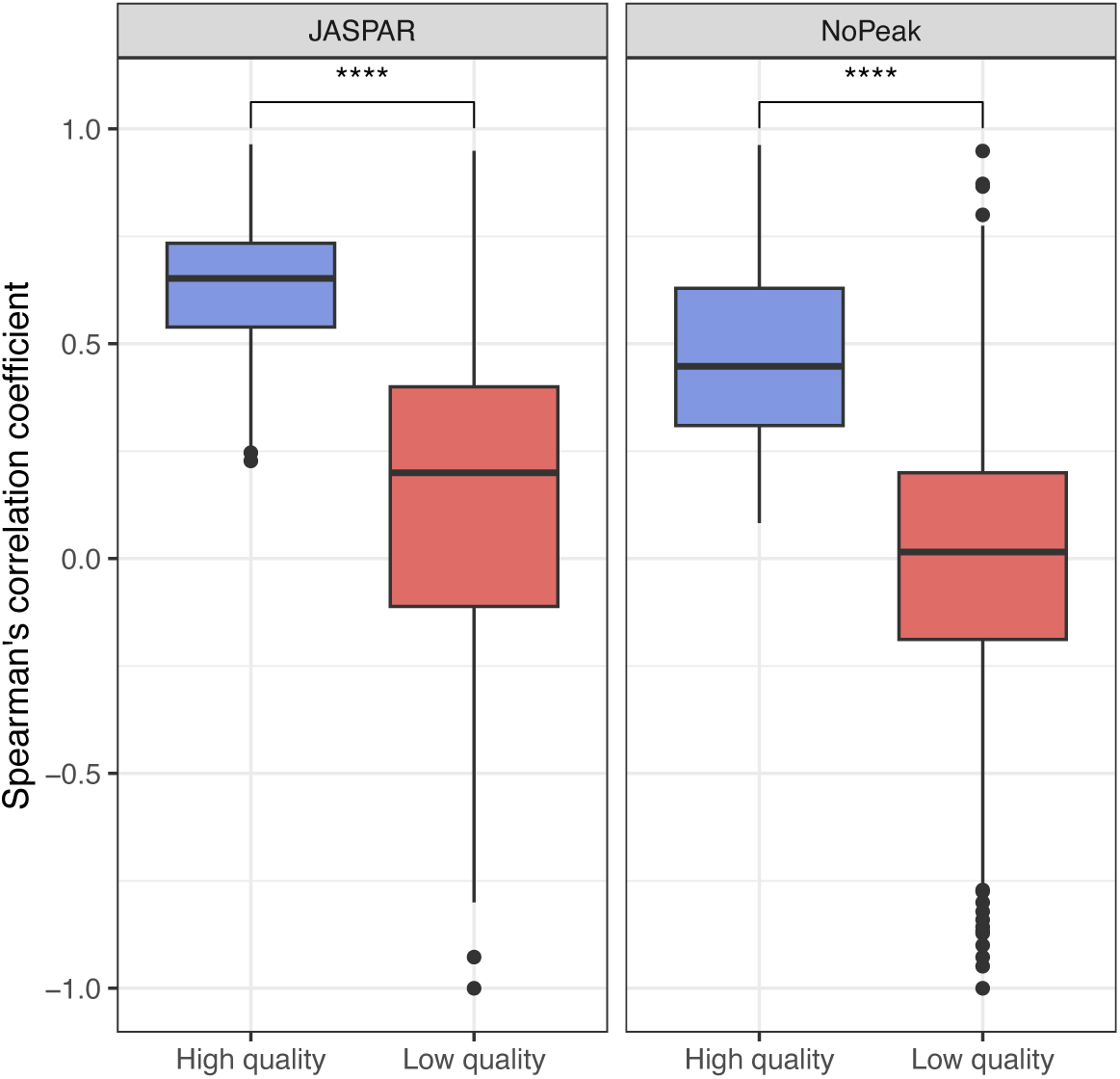
Spearman’s correlation coefficient for high-(blue) versus low-(red) quality motifs in JASPAR and NoPeak motifs investigated across all 558 TFs. **** indicate a p-value < 2 × 10^-16^ with p-values computed using a Wilcoxon rank sum test.

**Supplementary Figure S7:**
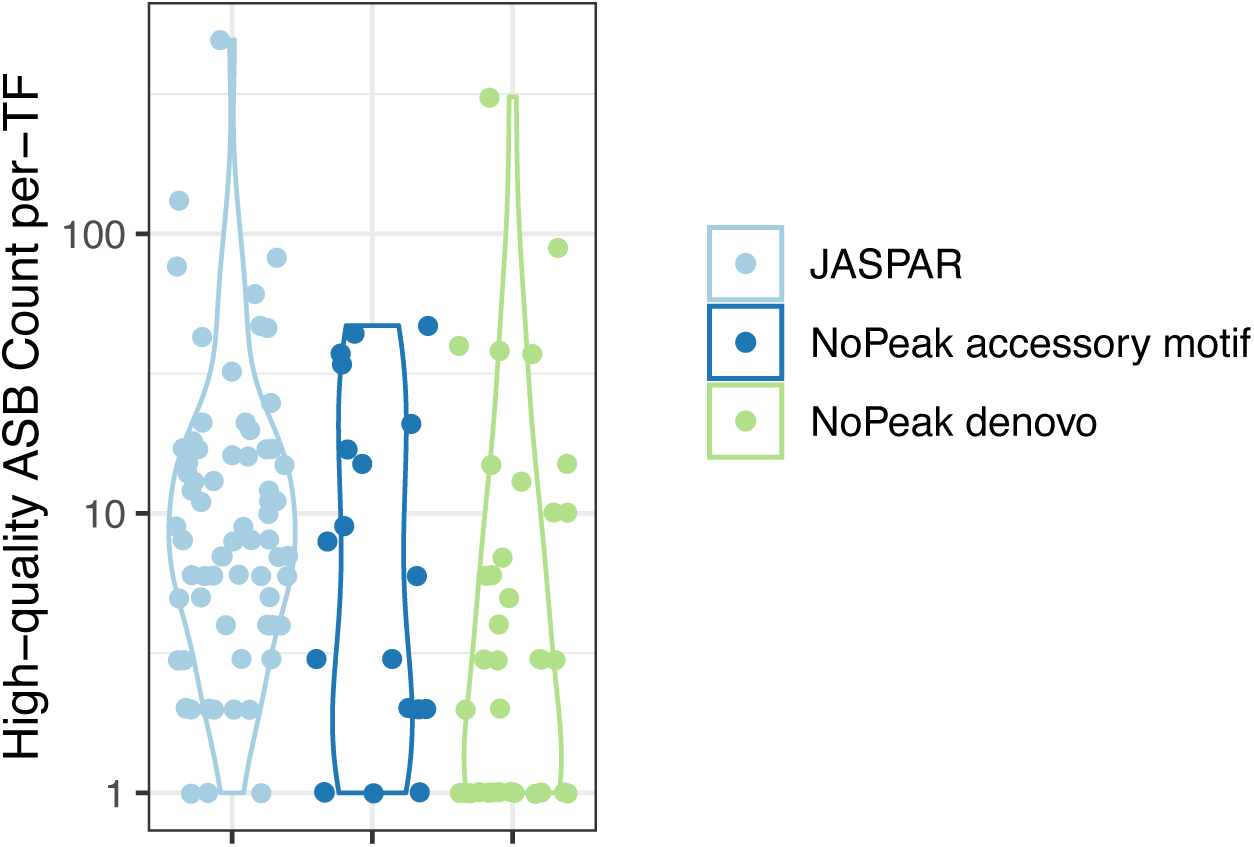
Breakdown by motif group for high-quality ASBs. Expanding our motif set to include accessory and de novo motifs from NoPeak substantially increases the number of high-quality ASBs.

**Supplementary Figure S8:**
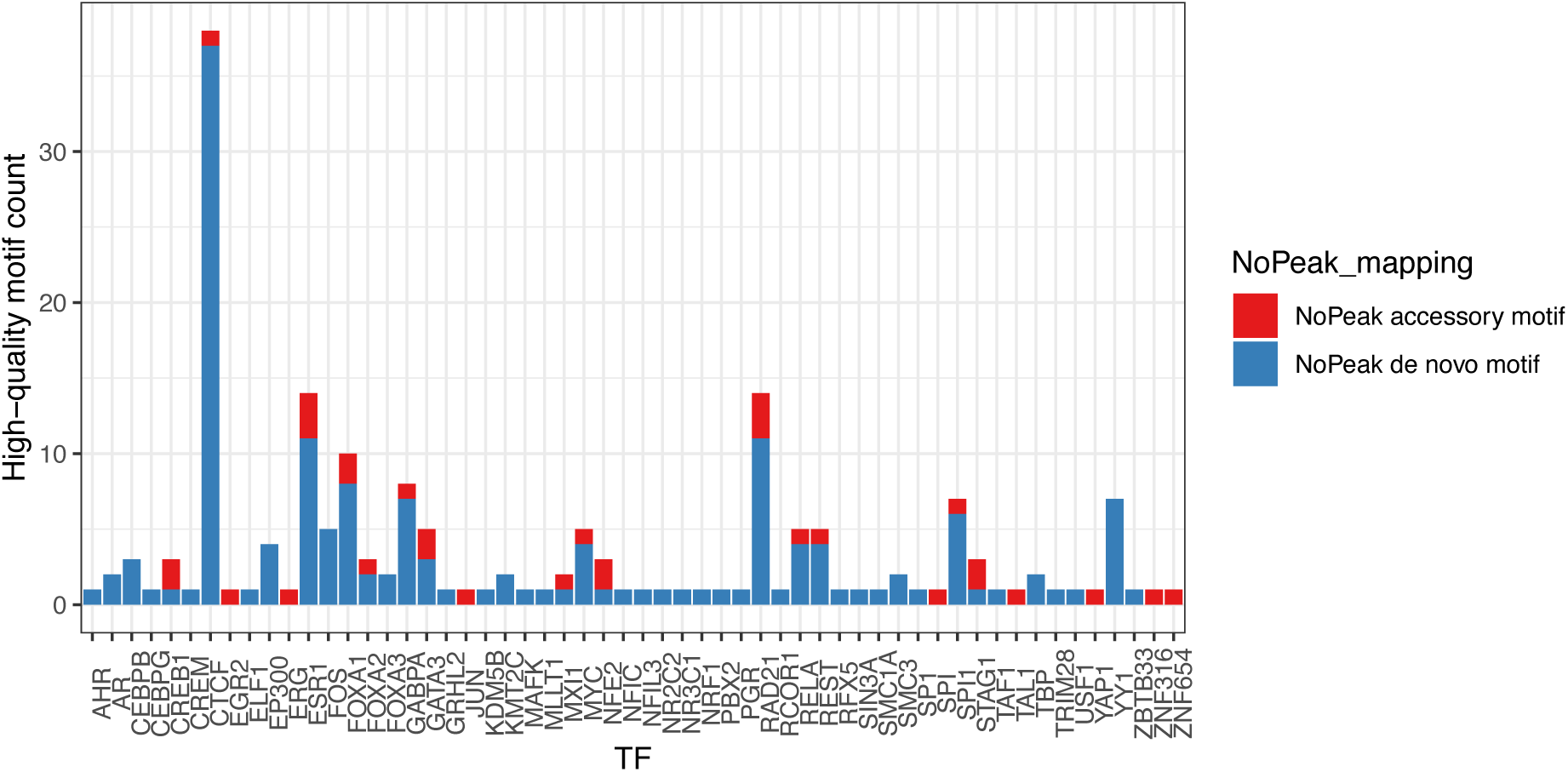
Numbers of NoPeak motifs detected across the 558 TF run. A range of NoPeak accessory and de novo motifs are discovered, with more de novo motifs than accessory motifs found overall.

**Supplementary Figure S9:**
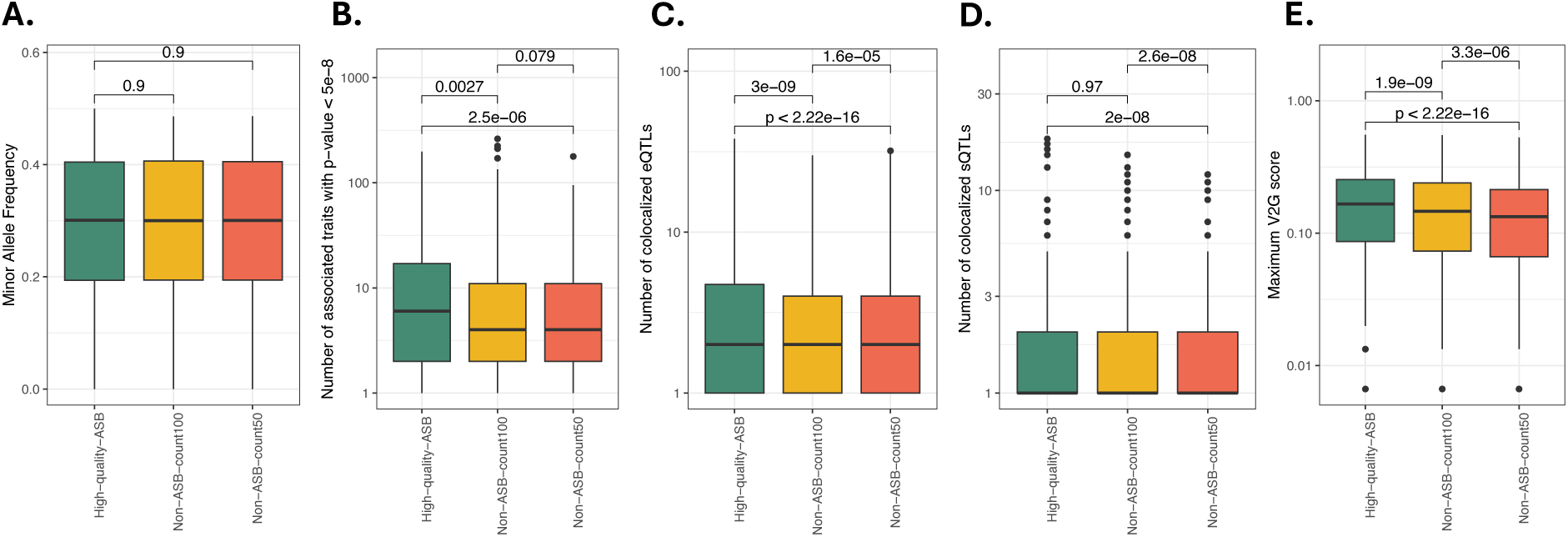
Trait/QTL associations for high-quality ASBs compared to non-ASB comparator sets. **(A)** As these non-ASB sets were matched on minor allele frequency (MAF), we see no significant difference between the MAFs for the high-quality ASB and non-ASB sets with p-values of 0.9 and 0.9 for the count 50 and count 100 thresholds, respectively. Non-ASB sets thresholded above 50 counts are enriched for **(B)** known variant-trait associations, and **(C)** colocalized eQTLs, but not for **(D)** sQTLs, although they tend to have **(E)** higher maximum V2G scores. Data is queried from OpenTargets for the median non-ASB set for counts threshold of 100 and 50 and the high-quality set. The median non-ASB set was defined by choosing the SNP with the median MAF across all 1000 sampled sets, for each SNP (see Methods). Note that the y-axis in plots **(B-E)** is log10-scaled.

**Supplementary Figure S10:**
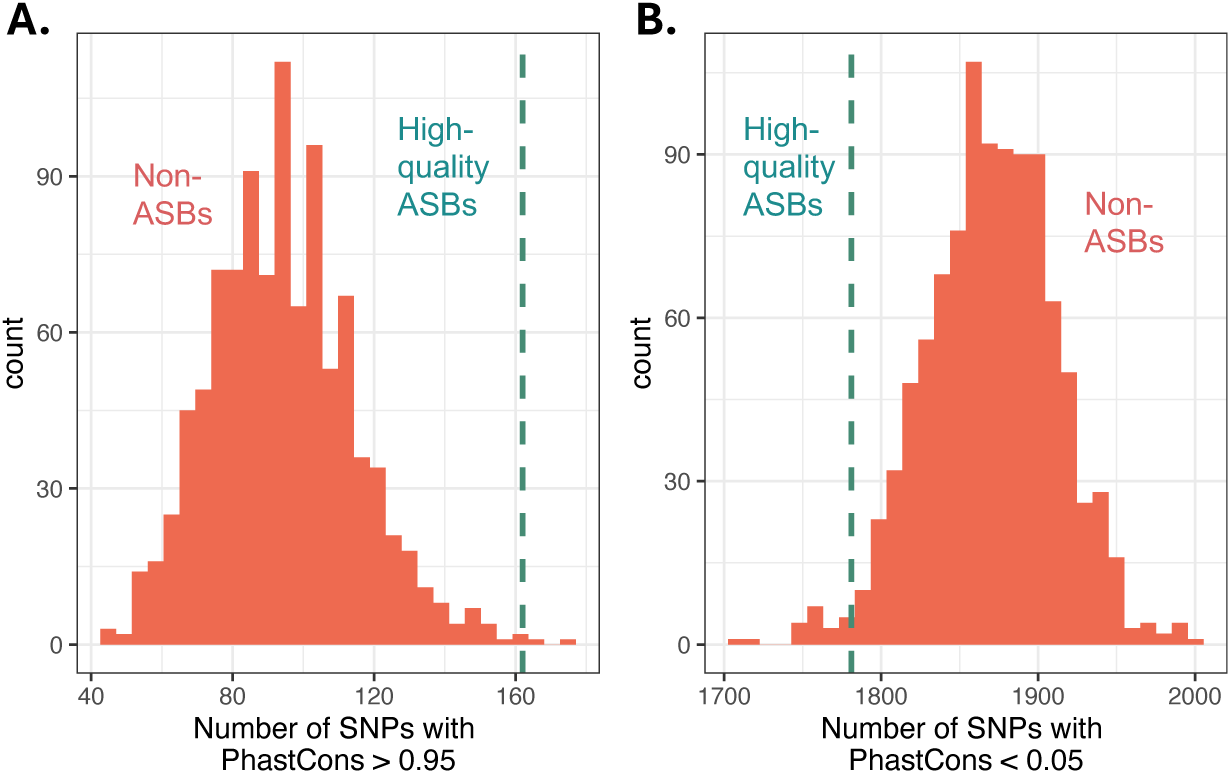
High-quality ASBs are more conserved than non-ASB sets across varying counts thresholds. This figure details the same analysis usng PhastCons scores shown in Figure 3 but for non-ASB sets with a minimum counts threshold of 50. **(A)** High-quality ASBs (dashed red line) harbour more conserved SNPs compared to count 50 non-ASB sets (green). **(B)** High-quality ASBs (dashed red line) have fewer non-conserved SNPs compared to count 50 non-ASB SNPs. PhastCons scores are pulled from the 30-way UCSC track for non-ASBs (green) and high-quality ASBs (red). The set of SNPs used to derive the sampled non-ASB sets was relaxed to a slightly lower read coverage minimum threshold of 50 counts. More specifically, the region containing the investigated SNP must not be called as an ASB in any cell line, for any TF, and have at least 50 reads mapping to it (see Methods). This is different to the analysis in the main figure which used a minimum count threshold of 100 reads.

**Supplementary Figure S11:**
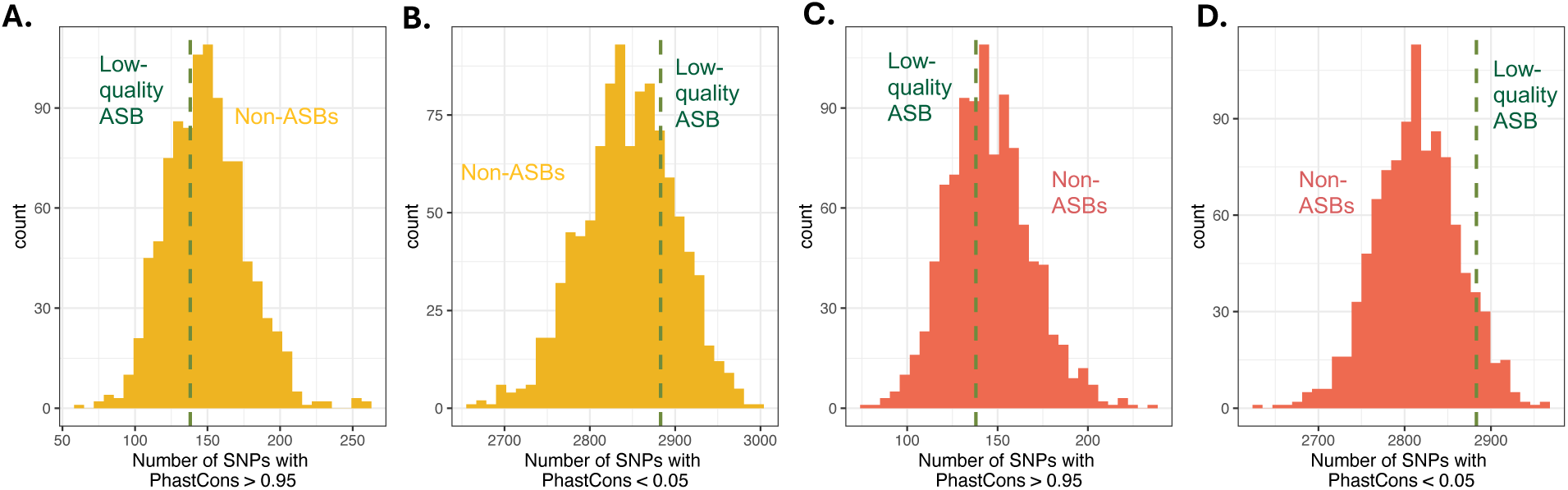
Low-quality ASBs are not better conserved than non-ASB sites. **(A)** No significant diference is found between the number of (A) highly-conserved SNPs or the number of **(B)** non-conserved SNPs when compared to the non-ASB set using a counts threshold > 100. Similar results are found for a counts threshold > 50 to determine non-ASB sets in **(C)** and **(D).**

**Supplementary Figure S12:**
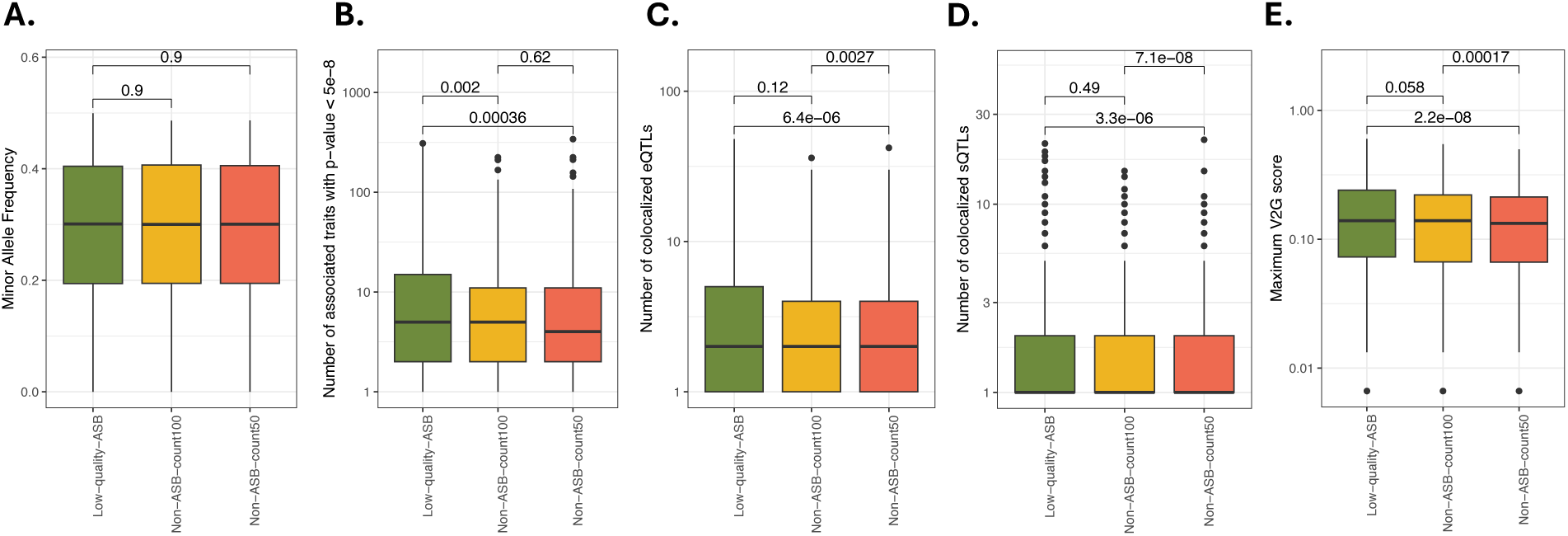
Trait/QTL associations for low-quality ASBs compared to non-ASB comparator sets. **(A)** As these non-ASB sets were matched on minor allele frequency (MAF), we see no significant difference between the MAFs for the low-quality ASB and non-ASB sets with p-values of 0.9 and 0.9 for the count 50 and count 100 thresholds, respectively. For low-quality ASBs, we find an enrichment for the **(B)** number of associated traits, but not for the number of colocalized **(C)** eQTLs, **(D)** sQTLs or **(E)** maximum V2G score. Data is queried from OpenTargets for the median non-ASB set for counts threshold of 100 and 50 and the low-quality set. The median non-ASB set was defined by choosing the SNP with the median MAF across all 1000 sampled sets, for each SNP (see Methods). Note that the y-axis in plots **(B-E)** is log10-scaled.

**Supplementary Figure S13:**
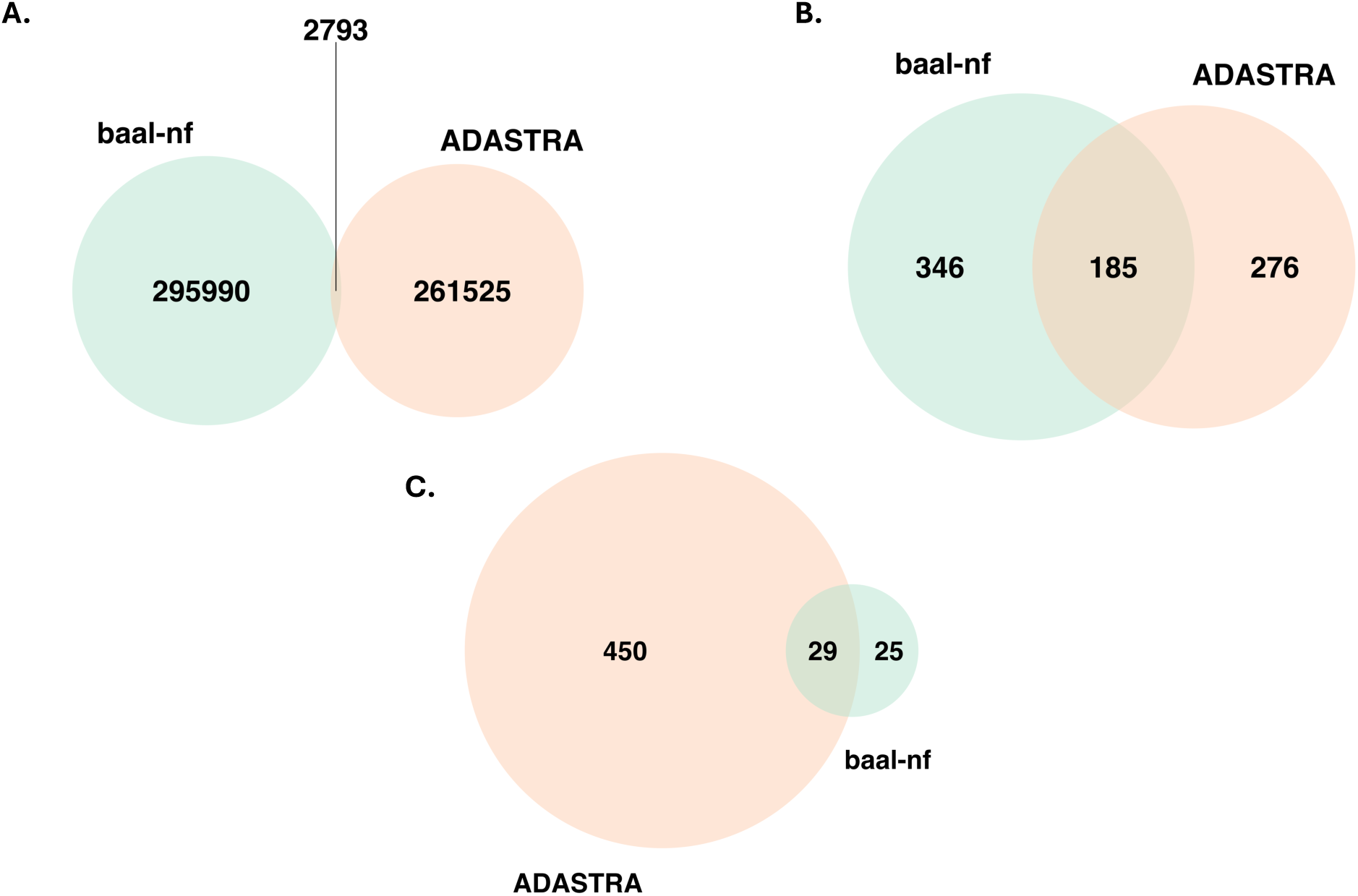
Overlap in features between ADASTRA ASB database and baal-nf ASB database. 295,990 ASBs (99%) predicted by baal-nf have not previously been reported by the ADASTRA resource [Abramov et al. (2021)] due largely to minimal overlap in the sets of TFs and cell lines investigated. Additional differences will be caused by ADASTRA not requiring heterozygous SNPs to be genotyped prior to its calling of ASBs. We find that **(A)** only 2,793 ASBs are found in both databases, meaning that 295,990 predicted by baal-nf (green) have not been previously reported by ADASTRA (orange). Notably, many of the features of tested SNPs were different across the two databases, including the **(B)** transcription factors investigated and **(C)** cell lines investigated.

**Supplementary Figure S14:**
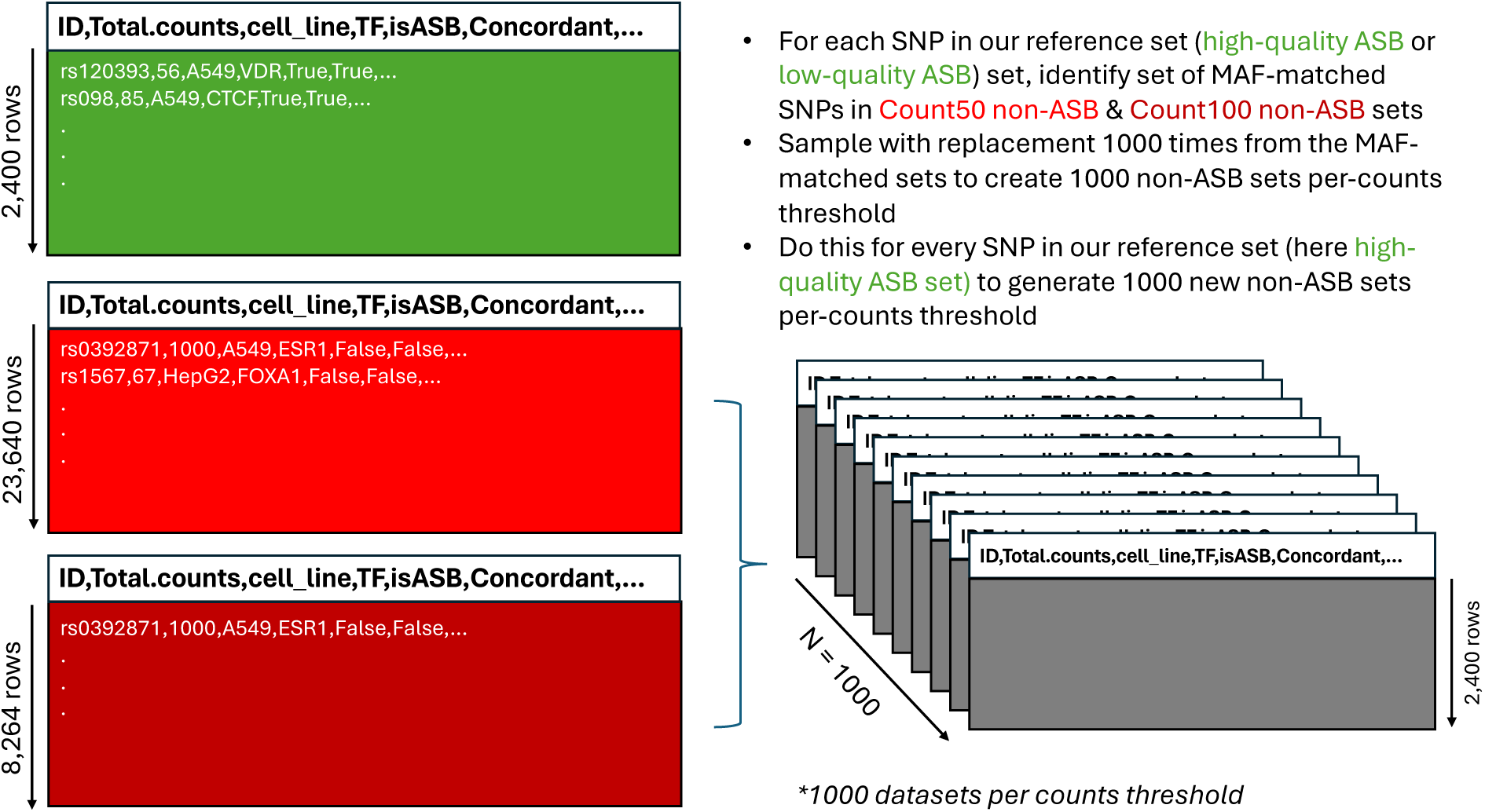
Flow chart detailing the sampling procedure for generating non-ASB sets from a reference set. (high-quality ASB set as reference shown here).

**Supplementary Figure S15:**
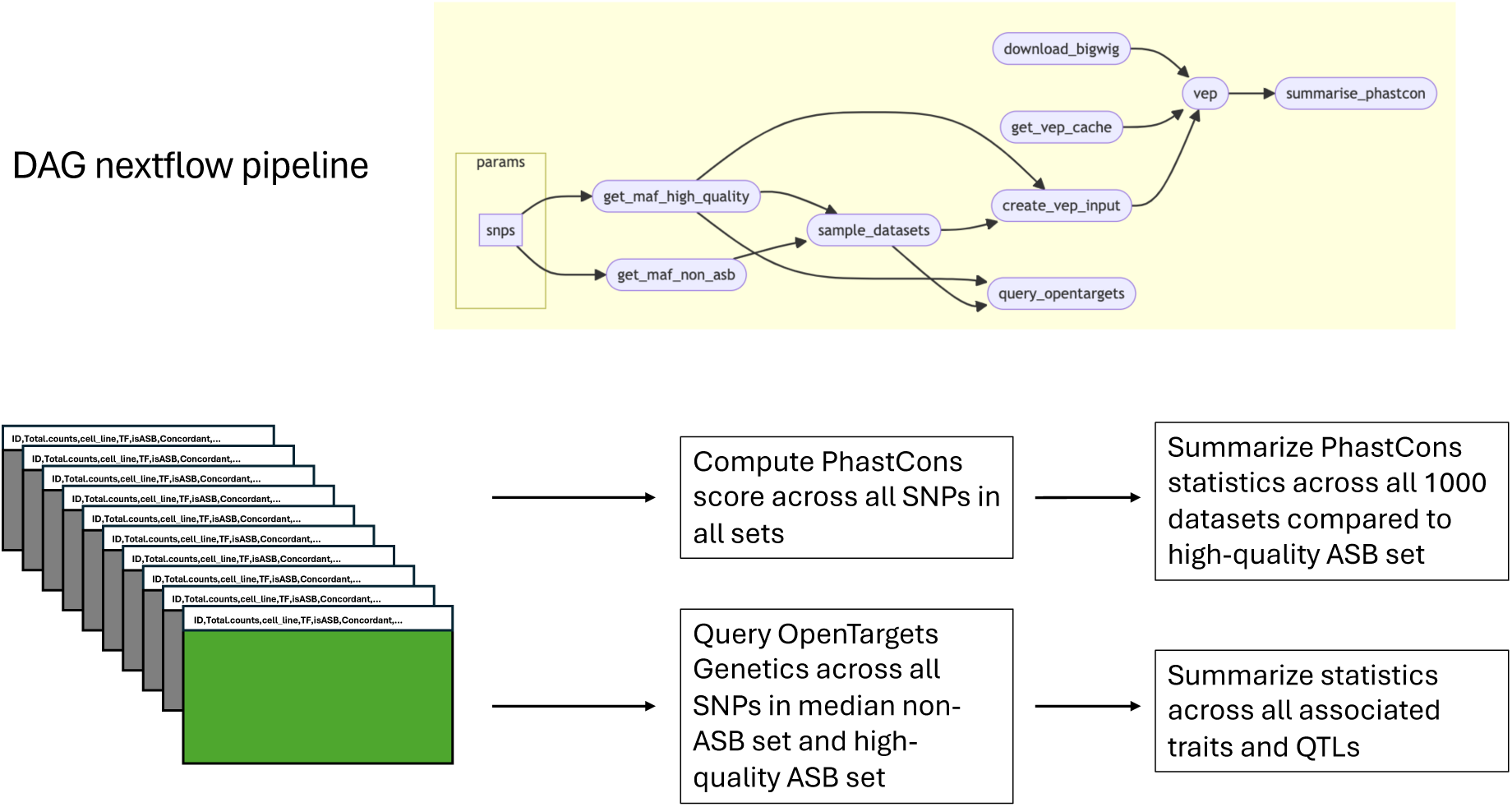
Pipeline for evaluating evolutionary, functional and trait relevance of non-ASB sets compared to reference ASB set. (shown here as high-quality ASB set).

## Notes

https://osf.io/rwjec/

https://github.com/BAAL-NF/baal-nf

https://www.encodeproject.org/

https://chip-atlas.org/

### Competing Interest Statement

The authors have declared no competing interest.

